# AMON: Annotation of metabolite origins via networks to better integrate microbiome and metabolome data

**DOI:** 10.1101/439240

**Authors:** M. Shaffer, K. Quinn, K. Doenges, X. Zhang, S. Bokatzian, N. Reisdorph, CA. Lozupone

## Abstract

**Motivation:** Untargeted metabolomics of host-associated samples has yielded insights into mechanisms by which microbes modulate health. However, data interpretation is challenged by the complexity of origins of the small molecules measured, which can come from the host, microbes that live with the host, or from other exposures such as diet or the environment.

**Results:** We address this challenge through development of AMON: Annotation of Metabolite Origins via Networks. AMON is an open-source bioinformatics application that can be used to determine the degree to which annotated compounds in the metabolome may have been produced by bacteria present, the host, either (i.e. both the bacteria and host are capable of production), or neither (i.e. neither the human or the fecal microbiome are predicted to be capable of producing the observed metabolite).

**Availability and Implementation:** This software is available at https://github.com/lozuponelab/AMON as well as via pip.

## INTRODUCTION

The host-associated microbiome can influence many aspects of human health and disease through its metabolic activity. Examples include microbe-host co-metabolism of dietary choline/carnitine to TMAO as a driver of heart disease (Wang *et al.*, 2011), microbial production of branched chain amino acids as a contributor to insulin resistance (Pedersen *et al.*, 2016), and microbial production of 12,13-DiHOME as a driver of CD4^+^ T cell dysfunction associated with childhood atopy (Fujimura *et al.*, 2016). A key way of exploring which compounds might mediate relationships between microbial activity and host disease is untargeted metabolomics (e.g. mass spectrometry) of host materials such as stool, plasma, urine, or tissues. These analyses result in the detection and relative quantitation of hundreds to thousands of compounds, the sum of which is referred to as a “metabolome”. Host-associated metabolomes represent a complex milieu of compounds that can have different origins, including the diet of the host organism and a variety of environmental exposures such as pollutants. In addition, the metabolome contains metabolic products of these compounds, i.e. metabolites, that can result from host and/or microbiome metabolism or co-metabolism (Shaffer *et al.*, 2017).

One way to estimate which metabolites in host samples originate from host versus microbial metabolism is to use metabolic networks described in databases such as KEGG (Kanehisa *et al.*, 2017). These networks capture the relationship between metabolites, the enzymes that produce them, and the genomes of organisms (both host and microbial) that contain genes encoding those enzymes. These networks provide a framework for relating the genes present in the host and colonizing bacteria, and the metabolites present in a sample.

Here we present AMON, which uses information in KEGG to predict whether measured metabolites are likely to originate from singular organisms or collections of organisms based on a list of the genes that they encode. As an example, AMON can be used to predict whether metabolites may originate from the host itself versus from host-associated microbiomes as assessed with 16S ribosomal RNA (rRNA) gene sequences or shotgun metagenomics. We demonstrate our tool by applying it to a dataset from a cohort of HIV positive individuals and controls in which the stool microbiome was assessed with 16S rRNA gene sequencing and the plasma metabolome was assessed with untargeted liquid chromatography mass spectrometry (LC/MS). We also illustrate how much information is lost when we only focus on compounds and genes of known identity/function, emphasizing the need for complimentary approaches to general metabolomic database searches for the identification of microbially produced compounds.

## METHODS

### AMON

AMON (Annotation of Metabolite Origins via Networks) is a command line tool for predicting which compounds are produced by microbes and which are produced by the host that is available at https://github.com/lozuponelab/AMON. The basis of this method is in multi-organism metabolic networks as depicted in Figure 1A. This is a directional network with a flow starting from nodes representing the organisms present in a community and edges connecting to the genes in the organism’s genomes. These genes connect to the chemical reactions that the proteins that they encode perform, which connect to the compounds that those reactions consume (incoming edges) and produce (outgoing edges). We trace up this network from compound to organism to determine the possible origin of a metabolite. For example, in Figure 1A, we can infer that the microbiome could have generated compound 9 because of the presence of gene 4 in the genome of Bacteria 2. However, compound 9 could also have been produced by the host because of the presence of gene 5 in the human genome. In contrast, compound 8 could only be produced by the bacteria present and not the host.

**Figure 1:**
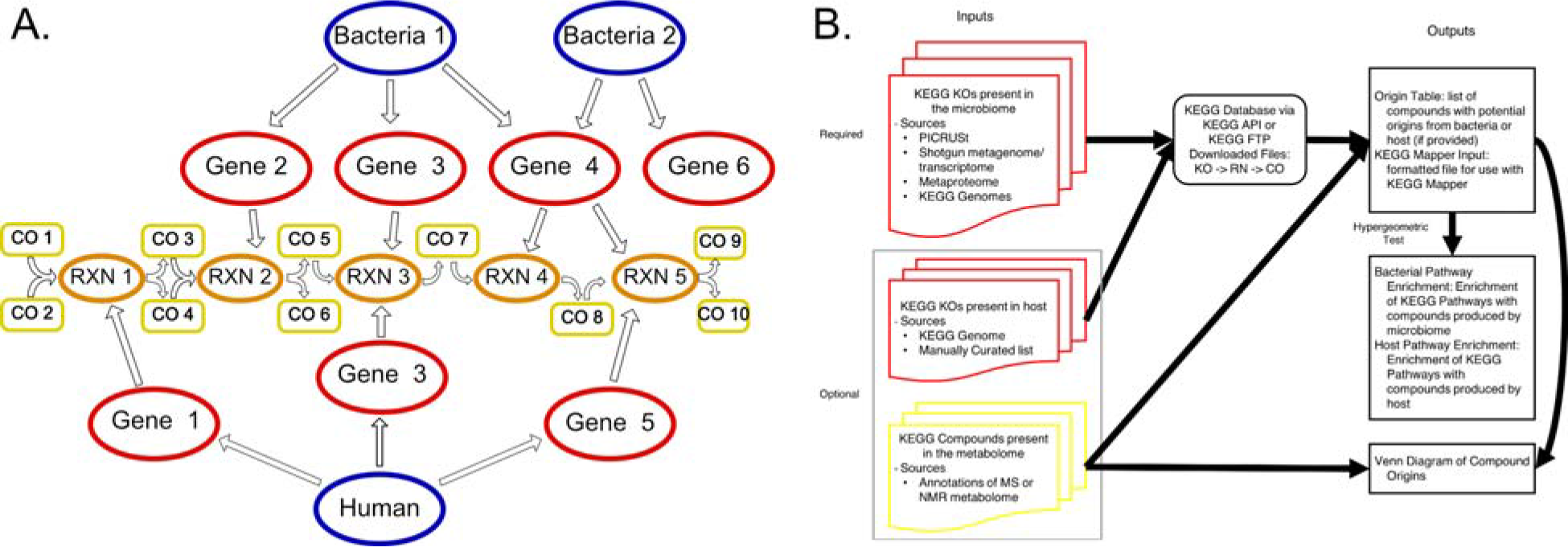
The network analysis and data flow of AMON. A) A simple multi-organism metabolic network. Blue nodes represent genomes, red nodes represent genes, orange nodes represent reactions and yellow nodes represent compounds. Edges between blue and red nodes indicate that the bacterial genomes contain the indicated genes and edges between red and orange nodes indicate the reactions which the genes can mediate. Yellow to orange edges connect reactants to a reaction and orange to yellow edges connect the reactions to its products. This network can be traversed to connect products of reactions to the genes and organisms which could produce these products. B) This schematic shows the flow of data through the AMON tool. The required input is a list of KEGG orthology (KO) identifiers which will be used with the KEGG database to build a metabolic network and determine the possible metabolites produced. This information is output to the user along with a pathway enrichment analysis to show functionality in the produced metabolite and a KEGG mapper file for visualization of metabolite origin in KEGG pathways.

AMON takes as input lists of KEGG KO (KEGG Orthology) identifiers that are predicted to be present in different potential sources (e.g. the metagenome of a host-associated microbiome or the genome of host organism) and a list of KEGG compound IDs, such as from an annotated metabolome (Figure 1B). AMON uses the multi-organism metabolic network constructed with information in KEGG to produce a table indicating which compounds (from the entire set of KEGG compounds and from the list of those annotated to be present in the metabolome) could be produced by each of the different provided KO sets and a file for input to KEGG mapper (https://www.genome.jp/kegg/mapper.html) which can be used to overlay this information on KEGG pathway diagrams. AMON uses the hypergeometric test to measure enrichment of KEGG pathways in metabolites predicted to originate specifically from each source environment that are present in the metabolome. Specifically, the set of metabolites predicted to be produced by the list of KO identifiers provided by the user is tested for enrichment of metabolites present in KEGG pathways relative to the background set of all compounds in all KEGG pathways that had at least one metabolite predicted to be produced by the provided gene sets. It produces a summary figure (Venn diagram) illustrating predicted metabolite origins.

AMON is built to be flexible as to the type of technology and informatics methods used to obtain the list of KOs present in each source sample and compounds present in a metabolome. As shown in our Case Study below, 16S rRNA data can be used to predict the KO list using PICRUSt (Langille *et al.*, 2013), which uses whole genome sequence information to predict KOs present. Other ways to produce this list of KOs include annotation of genes present in a shotgun metagenome, e.g. using tools such as HUMAnN (Abubucker *et al.*, 2012). The host KOs can be acquired from KEGG using the extract_ko_genome_from_organism.py script, which downloads the KOs from the KEGG API or parses them from a KEGG FTP file and makes a list of KOs present in that file.

AMON does not require the user to purchase a KEGG license. For individuals who have purchased a KEGG license, files containing KO and reaction information provided by KEGG can be loaded into AMON. As another option, AMON can also download the required information using the publicly available KEGG API (https://www.kegg.jp/kegg/rest/), although this method is comparatively slow and limits maximum dataset size based on the limits of the KEGG API.

### Case Study

We used AMON to relate the stool microbiome (as assessed with 16S rRNA gene sequencing) to the plasma metabolome (as assessed with untargeted LC/MS), in a cohort of HIV positive individuals (n=37) and HIV-negative controls (n=22). These data represent a subset of the cohort described in (Armstrong *et al.*, 2018) and are paired with metabolome data as a part of a study described at ClinicalTrials.gov (Identifier: NCT02258685). The overall goal of our case study was to use AMON to determine the degree to which annotated compounds in the plasma metabolome of our study cohort may have been produced by bacteria present in fecal samples, the host, either (i.e. both are capable of production), or neither (i.e. neither the human or the fecal microbiome are predicted to be capable of producing the observed metabolite).

All study participants were recruited from University of Colorado Hospital with an approved IRB protocol (CoMIRB 14-1595). Stool samples from 59 individuals were collected at home in a commode specimen collector within 24 hours of the clinic visit in which blood was drawn. Stool samples were stored at −20°C during transit and at −80°C prior to DNA extraction with the MoBIO kit and preparation for barcoding sequencing using the Earth Microbiome Project protocol (http://www.earthmicrobiome.org/protocols-and-standards/16s/). The 16S rRNA gene V4 region of stool microbes was sequenced using MiSeq (Illumina), denoised using DADA2 (Callahan *et al.*, 2016) and binned into 99% Operational Taxonomic Units (OTUs) using UCLUST (Edgar, 2010) and the greengenes database (version 13_8) via QIIME 1.9.1 (Caporaso *et al.*, 2010). We used PICRUSt (Langille *et al.*, 2013) to predict a metagenome and AMON to predict metabolites.

### Plasma Sample Preparation

A modified liquid-liquid extraction protocol was used to extract hydrophobic and hydrophilic compounds from the plasma samples (Yang *et al.*, 2013). Briefly, 100 μL of plasma spiked with internal standards underwent a protein crash with 400 μL ice cold methanol. The supernatant was dried under nitrogen and methyl *tert*-butyl ether (MTBE) and water were added to extract the hydrophobic and hydrophilic compounds, respectively. The upper hydrophobic layer was transferred to a new tube and the lower hydrophilic layer was re-extracted with MTBE. The upper hydrophobic layer was combined, dried under nitrogen and reconstituted in 200 μL of methanol. The hydrophilic layer was dried under nitrogen, underwent a second protein crash with water and ice-cold methanol (1:4 water-methanol). The supernatant was removed, dried by SpeedVac at 45 °C and reconstituted in 100 μL of 5% acetonitrile in water. Both fractions were stored at −80 °C until LCMS analysis.

### Liquid Chromatography Mass Spectrometry

The hydrophobic fractions were analyzed using reverse phase chromatography on an Agilent Technologies (Santa Clara, CA) 1290 ultra-high precision liquid chromatography (UHPLC) system on an Agilent Zorbax Rapid Resolution HD SB-C18, 1.8um (2.1 × 100mm) analytical column with an Agilent Zorbax SB-C18, 1.8 micron (2.1 × 5 mm) guard column. The hydrophilic fractions were analyzed using hydrophilic interaction liquid chromatography (HILIC) on a 1290 UHPLC system using a Phenomenex Kinetex HILIC, 2.6um (2.1 × 50mm) analytical column with an Agilent Zorbax Eclipse Plus C8 5μm (2.1 ×12.5mm) guard column. The hydrophobic and hydrophilic fractions were run on Agilent Technologies (Santa Clara, CA) 6520 and 6550 Quadrupole Time of Flight (QTOF) mass spectrometers, respectively. Both fractions were run in positive and negative electrospray ionization (ESI) modes, as previously described (Heischmann *et al.*, 2016).

### Mass Spectrometry Data Processing

Compound data was extracted using Agilent Technologies (Santa Clara, CA) Mass Hunter Profinder Version B.08 (Profinder) software in combination with Agilent Technologies Mass Profiler Professional Version 14 (MPP) as described previously (Heischmann *et al.*, 2016). Briefly, a naive feature finding algorithm, Find By Molecular Feature, was used in Profinder to extract compound data from all samples and sample preparation blanks. To reduce the presence of missing values, a theoretical mass and retention time database was generated for compounds present in samples only. This database was then used to re-search the raw sample data in Profinder using the Find By Ion algorithm.

An in-house database containing METLIN, Lipid Maps, KEGG, and HMDB spectral data was used to putatively annotate metabolites based on exact mass, isotope ratios and isotopic distribution with a mass error cutoff of 10 ppm. This corresponds to a Metabolomics Standards Initiative metabolite identification level 2 (Sumner *et al.*, 2007).

## RESULTS

We used PICRUSt to determine the genome content of the OTUs detected in the fecal samples. PICRUSt drops from the analysis OTUs that do not have related reference sequences in the database and produces an estimate of the nearest sequenced taxon index (NSTI) which measures how close those sequences are to sequenced genomes (those more closely related to genomes have more power to make predictions regarding gene content). Since human gut bacteria are well represented in genome databases, only 0.7% of total reads of the detected sequences were dropped on account of not having a related reference sequence in the database. Furthermore, the average NSTI across samples was 0.08, indicating that most OTUs were highly related to an organism with a sequenced genome. We applied PICRUSt to the 16S rRNA dataset with only OTUs present in more than 11 of 59 samples included. The 267 remaining OTUs were predicted to contain 4,409 unique KOs using PICRUSt. We used the KEGG list of KOs in the human genome to represent human gene content.

We provided these lists of gut microbiome and human KOs to AMON to produce a list of compounds generated from the gut microbiome and the human genome. Of the 4,409 unique KOs that PICRUSt predicted to be present in the gut microbiome, only 1,476 (33.5%) had an associated reaction in KEGG. Those without associated reactions may represent orthologous gene groups that do not perform metabolic reactions (such as transporters), or that are known to exist but for which the exact reaction is unknown, showing gaps in our knowledge (Fig 2A). Using information in KEGG, AMON predicted these KOs to produce 1,321 unique compounds via 1,926 unique reactions. The human genome was predicted to produce 1,376 metabolites via 1,809 reactions.

**Figure 2:**
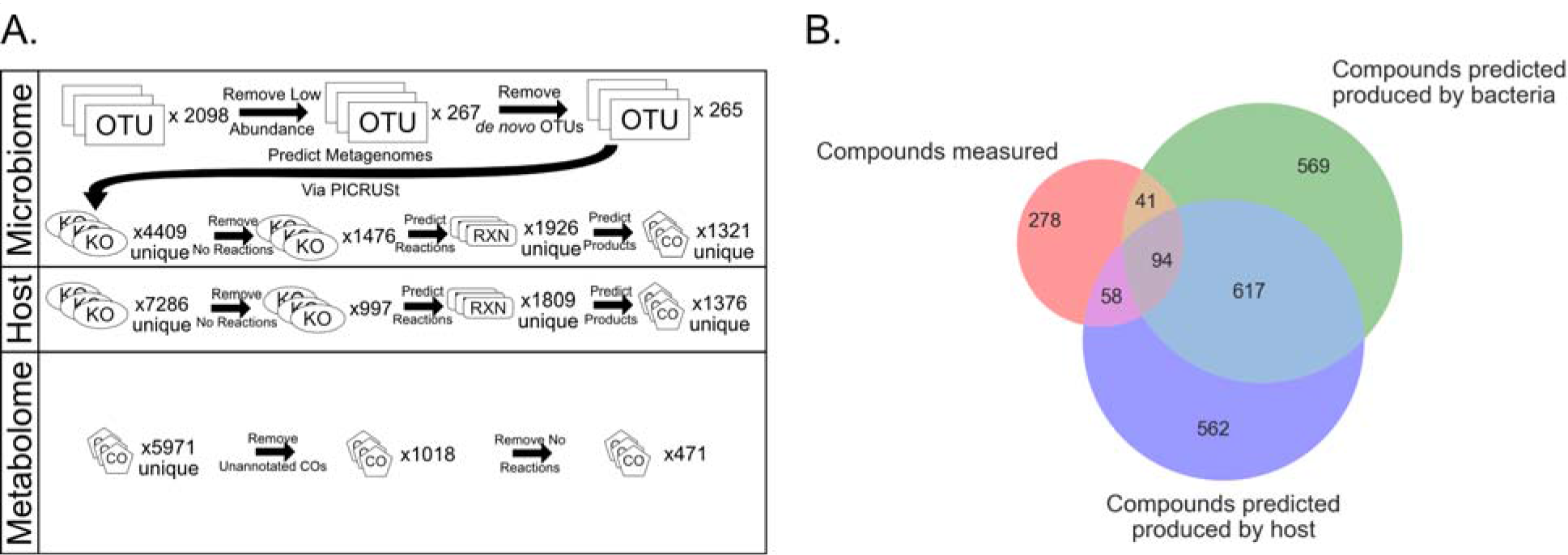
The results of a case study running AMON with 16S rRNA sequencing data from stool and PICRUSt to predict the metagenome along with the KEGG human genome and an LC/MS untargeted metabolome. A) A flow diagram showing how much data is lost between parts of analyses at all data levels. B) A Venn diagram showing overlaps in compound sets. The red circle shows compounds detected with untargeted LC/MS with an annotated KEGG compound ID. The green and purple circles show compounds that the metabolic network tells us could have been produced by the bacteria present in the microbiome and the host respectively.

Our metabolomics assays detected 5,971 compounds, of which only 1,018 (17%) could be putatively annotated with KEGG compound identifiers via a database search; only 471 (6%) of the 5,971 detected compounds were associated with a reaction in KEGG (Supplemental Table 1). Of these 471 annotated compounds in the plasma metabolome with associated KEGG reactions, 189 were predicted to be produced by enzymes in either human or stool bacterial genomes. 40 compounds were exclusively produced by bacteria, 58 exclusively by the host, and 91 by either human or bacterial enzymes (Fig 2B). The remaining 282 compounds may be 1) from the environment, 2) produced by microbes in other body sites or 3) host or gut microbial products from unannotated genes (Supplemental Table 1).

We used AMON to assess enrichment of pathways in the detected human and bacterial metabolites using the hypergeometric test (Figure 3A; Supplemental table 2). The 41 compounds predicted to be produced by stool bacteria and not the host were enriched in xenobiotic degradation pathways, including nitrotoluene and atrazine degradation, and pathways for amino acids metabolism, including the phenylalanine, tyrosine and tryptophan biosynthesis pathway and the cysteine and methionine metabolism pathway. The metabolite origin data was visualized using KEGG mapper for the phenylalanine, tyrosine and tryptophan biosynthesis pathway (Figure 3B). This tool helps to visualize the host-microbe co-metabolism and which genes are important for compounds that may have come from multiple sources. For instance, Figure 3B allows us to see that Indole is a compound found in our metabolome that could only have been produced by bacterial metabolism via the highlighted enzyme (K01695, tryptophan synthase).

**Figure 3:**
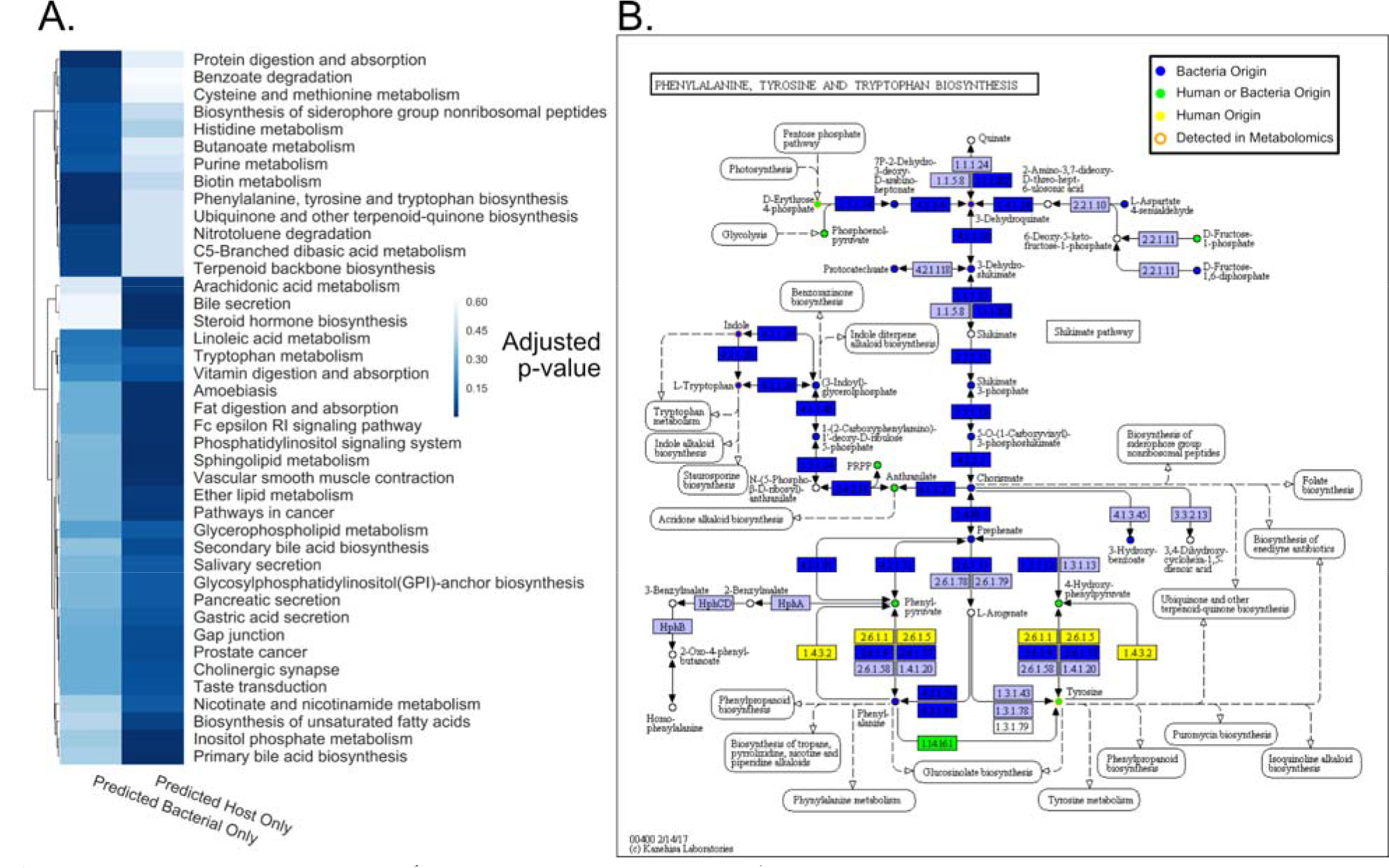
Enrichment of pathways and a single enriched pathway colored with metabolite origin. A) A heatmap showing the p-values associated with a pathway enrichment analysis with KEGG pathways. The first column is p-values for enrichment of KEGG pathways in compounds that were detected via untargeted LC/MS of plasma and we predict could be generated by members of the fecal microbiome. The second column is the same but for compounds that we predicted could have been generated by the human host. B) This pathway map is colored by putative origin of the compound, which are circles, and presence of the reaction, which are rectangles. Dark blue is a compound or gene with a bacterial origin, yellow is a compound or gene with a human origin, orange outlined compounds are detected in the metabolomics. Circles or rectangles could be of human or bacterial origin.

Also, Tyrosine is a compound found in our metabolome that could have been synthesized by a variety of enzymes found only in bacteria, only in humans, or in both and so further exploration would be needed to understand origins of this compound. The 51 compounds which were detected and predicted to be produced by the human genome were enriched in pathways that include bile secretion, steroid hormone biosynthesis and gastric acid secretion.

## DISCUSSION

Taken together, these analyses show that AMON can be used to predict the origin of compounds detected in a complex metabolome, such as stool. Our case study shows the specific application of predicting origins of plasma compounds as being from the fecal microbiome versus the host. However, this tool can be used to compare any number of different sources – e.g. from the microbiomes of different body sites or compounds that may come directly from plants consumed in the diet. Also, the outputs of AMON can be used in conjunction with lists of metabolites that were determined to significantly differ with disease state or correlate with other host phenotypes to predict origins of metabolites of interest.

Although our example uses PICRUSt to predict compounds of bacterial origin using 16S rRNA sequence data, AMON requires a list of KEGG Orthology identifiers as input and so could also be used with shotgun sequencing data. This can allow for a more thorough interrogation of host microbiomes that account for strain level variation in genome content and opens its application to environments with less understood genomes.

The pathway enrichment of compounds predicted to be unique to the gut microbiome and the host provide a level of validation for these results. The pathways enriched with compounds predicted to only be from microbes are consistent with known roles for gut bacteria in degrading various xenobiotics (Maurice *et al.*, 2013; Lu *et al.*, 2015; Das *et al.*, 2016; Saad *et al.*; Clayton *et al.*, 2009) and for influencing amino acid (Neis *et al.*, 2015; O’Mahony *et al.*, 2015) and vitamin metabolism (Streit and Entcheva, 2003). Likewise, the pathways enriched with compounds predicted to be human only include host processes such as taste transduction and bile secretion. Further, since the microbial community measured was from the human gut and the metabolome came from plasma, these results suggest that these microbial metabolites can translocate from the gut into systemic circulation. This is consistent with the gut microbiome being linked with many diseases that occur outside of the gut. Examples include interactions between the gut and brain via microbially derived compounds such as serotonin (O’Mahony *et al.*, 2015), and branched chain amino acids from the gut microbiome as a contributor to insulin resistance (Pedersen *et al.*, 2016).

However, this analysis also highlights limitations in this approach due to issues with annotation of both metabolites and the enzymes that may produce them. Overall, it is striking that of 5,971 compounds in the LC/MS data, only 471 could be linked to enzymatic reactions in KEGG. For example the human genome is known to contain approximately 20,000 genes (Ezkurdia *et al.*, 2014); however, there are only 7286 KOs annotated in KEGG. These KOs only predict the creation of 1376 unique compounds while the Human Metabolome Database 4.0 contains 114,100 (Wishart *et al.*, 2018). Part of this discrepancy is because multiple species of lipids are, generally, reduced to a single compound in KEGG. For example, while KEGG includes a single phosphatidylcholine (PC) lipid molecule in the Glycerophospholipid pathway, in fact, there are over 1,000 species of PCs. It is also important to note that metabolite annotations are based on peak masses and isotope ratios, which can often represent multiple compounds and/or in-source fragments; our confidence in the identity of these compounds is only moderate.

The situation is even worse for complex microbial communities, where even fewer genes are of known function. Because of these gaps in our knowledge of metabolite production, efforts to identify microbially produced metabolites that affect disease should also use methods that are agnostic to these knowledge-bases. These include techniques such as 1) identifying highly correlated microbes and metabolites to identify potential productive/consumptive relationships that can be further validated 2) molecular networking approaches which take advantage of tandem mass spectroscopy data to annotate compounds based on similarity to known compounds with related MS/MS profiles (Watrous *et al.*, 2012) or 3) coupling LC/MS runs with data from germ-free versus colonized animals (Wang *et al.*, 2011; Rothhammer *et al.*, 2016; Hsiao *et al.*, 2013) or antibiotic versus non-antibiotic treated humans (Tang *et al.*, 2013; Antunes *et al.*, 2011). Because AMON takes only KO identifiers and can pull database information from the KEGG API or user provided KEGG files, it will become increasingly useful as KEGG improves as well as other parts of the annotation process.

Although our application is specifically designed to work with the KEGG database, similar logic could be used for other databases such as MetaCyc (Caspi *et al.*, 2014). Our tool also does not apply methods such as gap-filling (Thiele *et al.*, 2014; Orth and Palsson, 2010) and metabolic modeling (Orth *et al.*, 2010; Mendes-Soares and Chia, 2017) in its estimates. The goal is not to produce precise measurements of the contributions of the microbiome and host to the abundance of a metabolite. Rather, AMON is designed to annotate metabolomics results to give the user an understanding of whether specific metabolites could have been produced directly by the host or its microbiomes. If a metabolite is identified by AMON as being of microbial origin and is associated with a phenotype, this result should motivate the researcher to perform follow up studies. These can include confirming the identity of the metabolite, via methods such as tandem mass spectrometry, and performing experiments to confirm the ability of microbes of interest to produce the metabolite.

AMON also does not account for co-metabolism between the host and microbes. An example of this is the production of TMAO from dietary choline. Our tool would list TMAO as a host compound and its precursor TMA as a microbiome derived compound but would not indicate that TMAO could overall not be produced from dietary substrates unless a microbiome was present. Further inspection of metabolic networks may be needed to decipher these co-metabolism relationships.

When researchers are seeking to integrate microbiome and metabolome data, identifying the origin of metabolites measured is an obvious route. AMON facilitates the annotation of metabolomics data by tagging compounds with their potential origin, either as bacteria or host. This allows researchers to develop hypotheses about the metabolic involvement of microbes in disease.

## ACKNOWLEDGEMENTS

This work was supported by National Institutes of Health R01 DK104047 and the associated metabolomics supplement by the National Institutes of Health Common fund and 4 T15 LM009451-10. High performance computing was supported by a cluster at the University of Colorado Boulder funded by National Institutes of Health 1S10OD012300.

